# Doublecortin compacts the microtubule lattice through a DC1-dependent mechanism

**DOI:** 10.64898/2026.11.14.766698

**Authors:** Regina Agulto, Julie Schlanz, Carter St. Clair, Him Shweta, Valerie Siahaan, Yusheng Shen, Lily B. Stine, Megan F. Hampton, Samuel E. Lacey, Marcus Braun, Zdenek Lansky, Yale E. Goldman, Ruxandra Dima, Richard J. McKenney, Kassandra M. Ori-McKenney

## Abstract

Doublecortin (DCX) is a microtubule-associated protein essential for neuronal migration during brain development, yet the molecular mechanisms underlying its function on the microtubule remain incompletely understood. Here we show that DCX compacts the microtubule lattice, with the DC1 domain serving as the main driver of lattice compaction. Using *in vitro* reconstitution, optical trapping, and molecular dynamics simulations, we demonstrate that although DCX preferentially binds compacted microtubule lattices, it can also bind expanded lattices and remodel them by inducing ∼2-3 Å compaction at interdimer interfaces. This compaction stiffens microtubules and inhibits kinesin-1 and αTAT1 binding. The DC1 domain is necessary for lattice compaction, whereas DC2 is dispensable for this function. Molecular dynamics simulations reveal that DC1 binding induces shearing and hinge motions that stabilize compacted conformations absent from apo tubulin, thereby reducing lattice heterogeneity. Our findings establish DCX as a bona fide microtubule remodeler and provide a molecular framework for understanding how DCX contributes to neuronal development through the control of microtubule lattice conformation.

## INTRODUCTION

Molecular movements dictate the architecture of all organisms. The development of the mammalian cerebral cortex requires the precise orchestration of neuronal migration, a process that depends on the microtubule cytoskeleton^1,2^. Doublecortin (DCX) is a microtubule-associated protein (MAP) that is essential for early brain development, and mutations in the DCX gene account for nearly a quarter of all cases of lissencephaly-spectrum brain malformations that arise from neuronal migration failure^3–5^. Elucidating the molecular mechanism of action of DCX is therefore essential for understanding both normal brain development and the pathogenesis of lissencephaly-spectrum disorders.

DCX interacts with microtubules through two conserved tandem doublecortin domains (DC1 and DC2), with the DC1 domain playing a dominant role in microtubule binding^6–8^. The DC domains bind between protofilaments at interdimer interfaces, a site that changes conformation during GTP hydrolysis^6,9,10^. Consistent with this binding location, DCX specifically binds straight microtubules and is excluded from the GTP/GDP-P_i_ cap recognized by end-binding proteins (EBs)^11–15^. Prior work has shown that DCX and its orthologs may act as microtubule stabilizers, nucleators, and regulators of microtubule-based transport^14,16–18^. DCX binds microtubules cooperatively^11^, and this cooperativity likely underlies its ability to perform this myriad of functions. Recent studies have revealed that DCX is required for growth cone advance in soft microenvironments that mimic brain tissue viscoelasticity, where it contributes to growth cone biomechanics by reinforcing the cytoskeleton against actomyosin-generated contractile forces^15^. Additionally, DCX has been shown to restrict neuronal branching by regulating α-tubulin polyglutamylation^19^, and to regulate dynein-mediated retrograde transport through interactions with the dynein motor complex^20^. The molecular mechanism that connects these diverse cellular functions remains elusive.

Microtubules exist in distinct conformational states defined by the longitudinal lattice spacing of αβ-tubulin dimers that compose individual protofilaments. GTP-hydrolysis drives a conformational change from an expanded state (∼83 Å), associated with newly incorporated GTP-tubulin, to a compacted state characteristic of the GDP-lattice (∼81 Å)^10^. This lattice compaction is functionally significant as microtubule-associated proteins such as tau and MAP2 have been shown to bind and compact the microtubule lattice through cooperative protein “envelope” formation^21–23^. Conversely, other MAPs such as MAP7, MAP4, and CAMPSAP3 have been identified as microtubule expanders that stabilize the extended lattice conformation^22,24,25^, and small molecule drugs such as paclitaxel similarly promote lattice expansion^10,26^. Recent evidence has suggested that DCX may act as a compactor by competing with paclitaxel^26^, yet direct demonstration of DCX-induced lattice compaction and the molecular determinants of this activity are lacking.

Here, we investigate the molecular mechanism by which DCX interacts with and modulates microtubule lattice conformation. We find that DCX preferentially binds compacted lattices but can also bind expanded lattices and compact them. Using a combination of *in vitro* reconstitution, optical trapping, cryo-electron microscopy, and molecular dynamics simulations, we demonstrate that DC1 is the main driver of lattice compaction, while DC2 is dispensable for this function. DCX-induced lattice compaction stiffens microtubules and allosterically inhibits kinesin-1 motility, revealing a mechanism by which DCX may regulate both microtubule mechanics and motor access. Our findings establish DCX as a bona fide microtubule compactor, joining tau and MAP2 in a convergent evolutionary strategy to regulate microtubule conformation.

## RESULTS

### DCX Preferentially Binds to Compacted or Compactible Microtubule Lattices

To investigate the relationship between microtubule lattice conformation and the binding of doublecortin (DCX), we employed total internal reflection fluorescence microscopy (TIRF-M) using different microtubule lattices. Taxol-stabilized microtubules exist in a reversibly expanded conformation that can be compacted by effectors such as tau and MAP2, while GMPCPP microtubules are in an irreversibly expanded state^10,21,22^. Native GDP microtubules represent the compacted conformation that results from GTP hydrolysis following tubulin incorporation into the lattice^10,27^.

We first compared the binding of DCX to taxol-stabilized and GMPCPP microtubules. At 20 nM, DCX exhibited significantly higher fluorescence intensity on taxol-stabilized microtubules compared to GMPCPP microtubules (Figure 1A-B). This indicates that although DCX can bind both lattice types, it preferentially associates with taxol-stabilized microtubules, which can undergo compaction upon DCX binding. Alternatively, or in addition, the increased DCX intensity may indicate that the compacted lattice accommodates a higher density of DCX binding. To directly test DCX’s preference for the native compacted lattice, we polymerized native GDP microtubules from GMPCPP seeds, creating a hybrid structure where the seed remains expanded and the newly polymerized lattice is compacted. DCX showed markedly increased binding to the GDP lattice compared to the adjacent GMPCPP lattices (Figure 1C-E). These results are consistent with prior work showing DCX exhibits reduced binding to GMPCPP-stabilized lattices compared to dynamic microtubules^11^ and demonstrates that DCX preferentially binds to compacted or compactible microtubule lattices.

**Figure 1.**
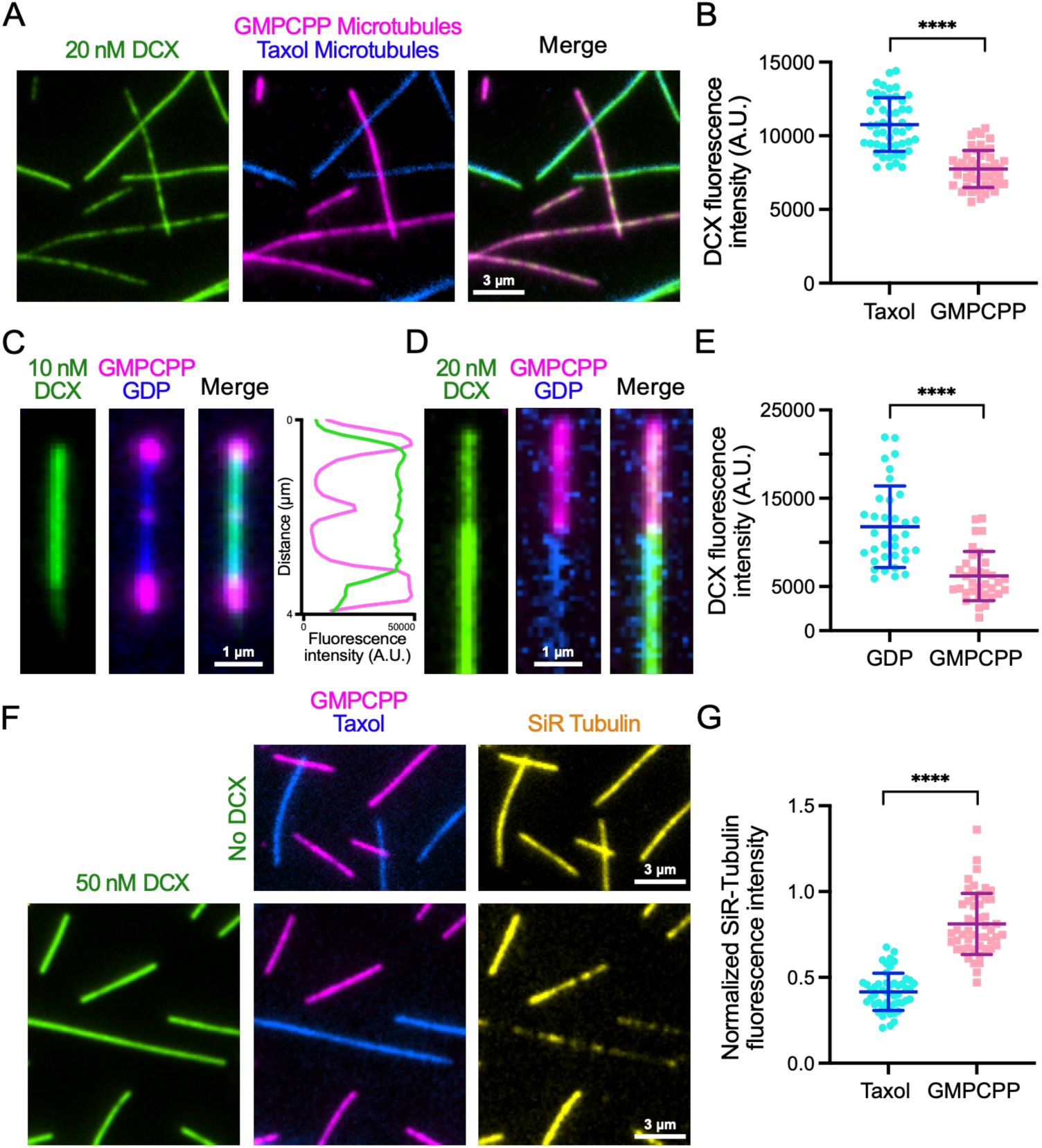
DCX Preferentially Binds to Compacted or Compactible Microtubule Lattices. **(A)** Total internal reflection fluorescence microscopy (TIRF-M) images of 20 nM sfGFP-DCX (green) binding to taxol-stabilized microtubules (blue) and GMPCPP microtubules (magenta). Scale bars = 3 µm. **(B)** Quantification of DCX fluorescence intensity on taxol-stabilized microtubules compared to GMPCPP microtubules. Means ± s.d.: 10763.8 ± 1818.6 for taxol-stabilized microtubules and 7744.7 ± 1257.7 for GMPCPP microtubules. *n* = 50 microtubules per condition from *n* = 2 independent trials. **(C)** TIRF-M images of 10 nM sfGFP-DCX (green) binding to a native GDP microtubule lattice (blue) polymerized from and capped by GMPCPP seeds (magenta). Associated line scan reveals increased fluorescence intensity of sfGFP-DCX on the GDP lattice compared with the GMPCPP lattices. Scale bar = 1 µm. **(D)** TIRF-M images of 20 nM sfGFP-DCX (green) binding to a native GDP microtubule lattice (blue) polymerized from a GMPCPP seed (magenta). Scale bar = 1 µm. **(E)** Quantification of 20 nM DCX fluorescence intensity on native GDP microtubule lattices compared to adjacent GMPCPP seeds. Fluorescence intensity is measured as background-subtracted mean intensity per unit length of microtubule. Means ± s.d.: 5546.1 ± 793.8 for GDP lattices and 4177.3 ± 1270.6 for GMPCPP lattices. *n* = 50 microtubules per condition from *n* = 2 independent trials. **(F)** TIRF-M images showing 1 µM SiR-Tubulin (yellow) binding to taxol-stabilized (blue) and GMPCPP microtubules (magenta) in the absence (top row) or presence (bottom row) of 50 nM sfGFP-DCX. Scale bars = 3 µm. **(G)** Quantification of SiR-Tubulin fluorescence intensity on taxol-stabilized microtubules compared to GMPCPP microtubules in the presence of 50 nM DCX normalized to SiR-Tubulin fluorescence intensity in the absence of sfGFP-DCX on each microtubule lattice. Means ± s.d.: 0.42 ± 0.11 for taxol-stabilized microtubules and 0.81 ± 0.18 for GMPCPP microtubules. *n* = 50 microtubules per condition from *n* = 2 independent trials. All graphs display all datapoints with lines indicating means ± s.d. **** indicates *p* < 0.0001 using two-tailed unpaired Student’s t-test.

To determine whether DCX compacts the microtubule lattice upon binding, we utilized SiR-tubulin, a docetaxel derivative that preferentially binds to expanded lattices^22,24,26^. We measured SiR-tubulin fluorescence intensity on taxol-stabilized and GMPCPP microtubules in the presence or absence of 50 nM DCX (Figure 1F). In the absence of DCX, SiR-tubulin intensity was comparable between lattice types. However, upon DCX binding, SiR-tubulin intensity on taxol-stabilized microtubules decreased by approximately 58% compared to untreated controls, while the intensity on GMPCPP microtubules decreased 19% (Figure 1G). This differential reduction in SiR-tubulin association indicates that DCX compacts the taxol-stabilized lattice upon binding, displacing the expanded-lattice probe. The minimal change on GMPCPP microtubules confirms that DCX is unable to efficiently compact lattices locked in the expanded state. Thus, DCX binds to both expanded and compacted lattices, but exhibits higher occupancy on compacted lattices and compacts expanded lattices upon binding.

### DCX Compacts the Microtubule Lattice

DCX is known to bind microtubules cooperatively, which contributes to the formation of dense DCX clusters on the microtubule surface^11^. Similar cooperative assemblies, termed “envelopes,” have been described for tau and MAP2, which form cohesive structures that compact the microtubule lattice and regulate the accessibility of the lattice by other MAPs and motors^21,22^. Consistent with these observations, we previously showed that DCX clusters act as selectively permeable barriers that regulate motor and MAP access to the microtubule surface^18^. Based on the similarity between DCX clustering behavior and tau/MAP2 envelope formation, we next investigated whether DCX, like tau and MAP2, actively compacts the microtubule lattice. We employed three complementary approaches: optical trap-based force measurements, cryo-electron microscopy (cryo-EM), and speckle-tubulin imaging. In the optical trap assay, we suspended taxol-stabilized, biotinylated microtubules between two streptavidin-functionalized beads held in optical traps and measured the force-extension curve by repeatedly stretching the MT up to ∼25 pN of force and releasing it back to slack length. The bead-to-bead distance at 25 pN tension decreased upon moving the microtubule into a channel containing 50 nM DCX (Figure 2A), indicating microtubule lattice compaction (Figure 2B). In contrast, when we performed the identical experiment with 50 nM MAP4, a microtubule-associated protein that recognizes expanded lattices without compacting them^22^, no such decrease in bead-to-bead distance was observed (Figure 2C-D). Strikingly, DCX compacted the lattice by 2.38% (Figure 2E), which closely matches the length difference between GTP- and GDP-bound tubulin dimers estimated previously by cryo-EM to be 2.4%^10^, suggesting DCX binding shifts the microtubule from its extended GTP-like state into a compacted GDP-like conformation.

**Figure 2.**
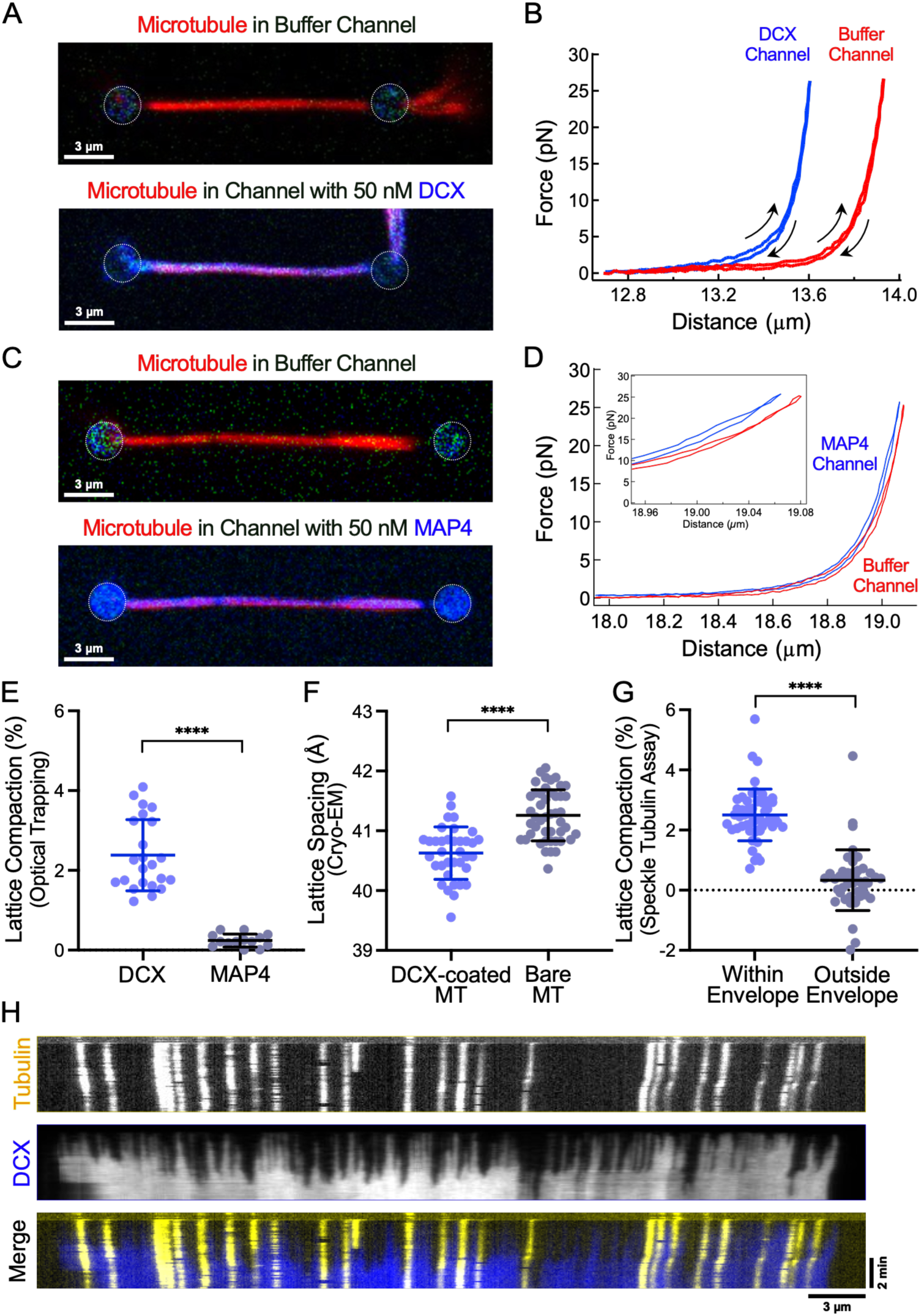
DCX Directly Compacts the Microtubule Lattice. **(A)** Confocal fluorescence images of a taxol-stabilized, biotinylated microtubule (red) suspended between two optically-trapped streptavidin-functionalized beads in the buffer channel (top), then moved into a channel containing 50 nM sfGFP-DCX (blue) (bottom). The beads are indicated by dashed white circles. Scale bars = 3 µm. **(B)** Force-extension curves of the microtubule in the buffer (red) and DCX (blue) channels. The stretch-relax cycles are indicated by up-arrows and down-arrows. **(C)** Confocal fluorescence images of a taxol-stabilized, biotinylated microtubule (red) suspended between two optically-trapped streptavidin-functionalized beads in the buffer channel (top), then moved into a channel containing 50 nM sfGFP-MAP4 (blue) (bottom). The beads are indicated by dashed white circles. Scale bars = 3 µm. **(D)** Force-extension curves of the microtubule in the buffer (red) and MAP4 (blue) channels. Binding of DCX induces a decrease in the distance between the two beads indicating microtubule compaction, whereas no such shift is detected in the MAP4 channel (see inset). **(E)** Quantification of percent compaction of microtubules upon the binding of DCX or MAP4. Means ± s.d. are 2.38 ± 0.89 from n = 23 dumbbells and 0.24 ± 0.16 from n = 14 dumbbells from several independent experiments for DCX and MAP4, respectively. **(F)** Quantification of the tubulin monomer spacing from cryo-EM images of taxol-lattice microtubules in the presence or absence of DCX. The bare microtubule control data (41.3 ± 0.4 Å from n = 46 microtubules) were previously reported^22^. In the presence of DCX, the spacing decreased to 40.6 ± 0.4 Å from n = 30 microtubules. Means ± s.d. are shown. **(G)** Quantification of microtubule lattice compaction by DCX inside envelope regions vs. outside envelope regions in a speckle-tubulin assay. Means ± s.d. are 2.5 ± 0.9 % from n = 52 microtubules and 0.3 ± 1.0 % from 44 microtubules from n = 5 independent experiments. **(H)** Representative kymograph from a speckle-tubulin assay showing microtubule lattice compaction as evidenced by the decreased distance over 5 min between individual tubulin speckles (yellow) with the addition of 30 nM sfGFP-DCX (blue). Scalebars: x = 3 µm, y = 2 min. All graphs display all datapoints with lines indicating means ± s.d. **** indicates *p* < 0.0001 using two-tailed unpaired Student’s t-test.

To quantify this effect at the molecular level, we performed cryo-EM analysis of taxol-stabilized microtubules in the presence or absence of DCX to directly measure the tubulin monomer spacing within the lattice. We found that DCX binding resulted in a significant decrease in the average spacing between tubulin monomers, from 41.3 ± 0.4 Å in the absence of DCX to 40.6 ± 0.4 Å in the presence of DCX (Figure 2F). This compaction brings the lattice spacing to that characteristic of the native GDP state.

Finally, to study the role of the cooperative binding mode of DCX in microtubule lattice compaction, we used a speckle-tubulin assay to measure compaction on individual microtubules partially covered by DCX envelopes. We added 30 nM DCX to taxol-stabilized microtubules containing fluorescent tubulin speckles and tracked speckle positions over time (Figure 2G-H)^22^. We observed minimal compaction of the lattice in microtubule regions not covered by DCX envelopes (0.3 %), while the lattice regions that were covered by DCX showed a compaction of 2.5% (Figure 2G). Collectively, these three independent approaches demonstrate that DCX directly compacts the microtubule lattice upon binding.

### The DC1 Domain Is Necessary for DCX-Induced Lattice Compaction

Having established that DCX directly compacts the microtubule lattice, we next sought to identify which regions of DCX are necessary for this function. Figure 3A shows block diagrams of a panel of purified DCX constructs including full-length WT, ΔDC1 (lacking the first DC domain), ΔDC2 (lacking the second DC domain), ΔC (lacking an intrinsically disordered C-terminal region), and various modifications of the 40 aa segment linking the two DC domains, including a GAS-40 linker (substituting the linker sequence with <u>g</u>lycine, <u>a</u>lanine, and <u>s</u>erine residues), a Scrambled linker (scrambling the wild-type aa sequence), a ½ Linker (deleting the last twenty amino acids of the linker, residues 160-179), and a 2X Linker (duplicating the linker region). The linker region length and pI is conserved among doublecortin family members (Figure S4), prompting us to systematically investigate its role in DCX-mediated lattice compaction.

**Figure 3.**
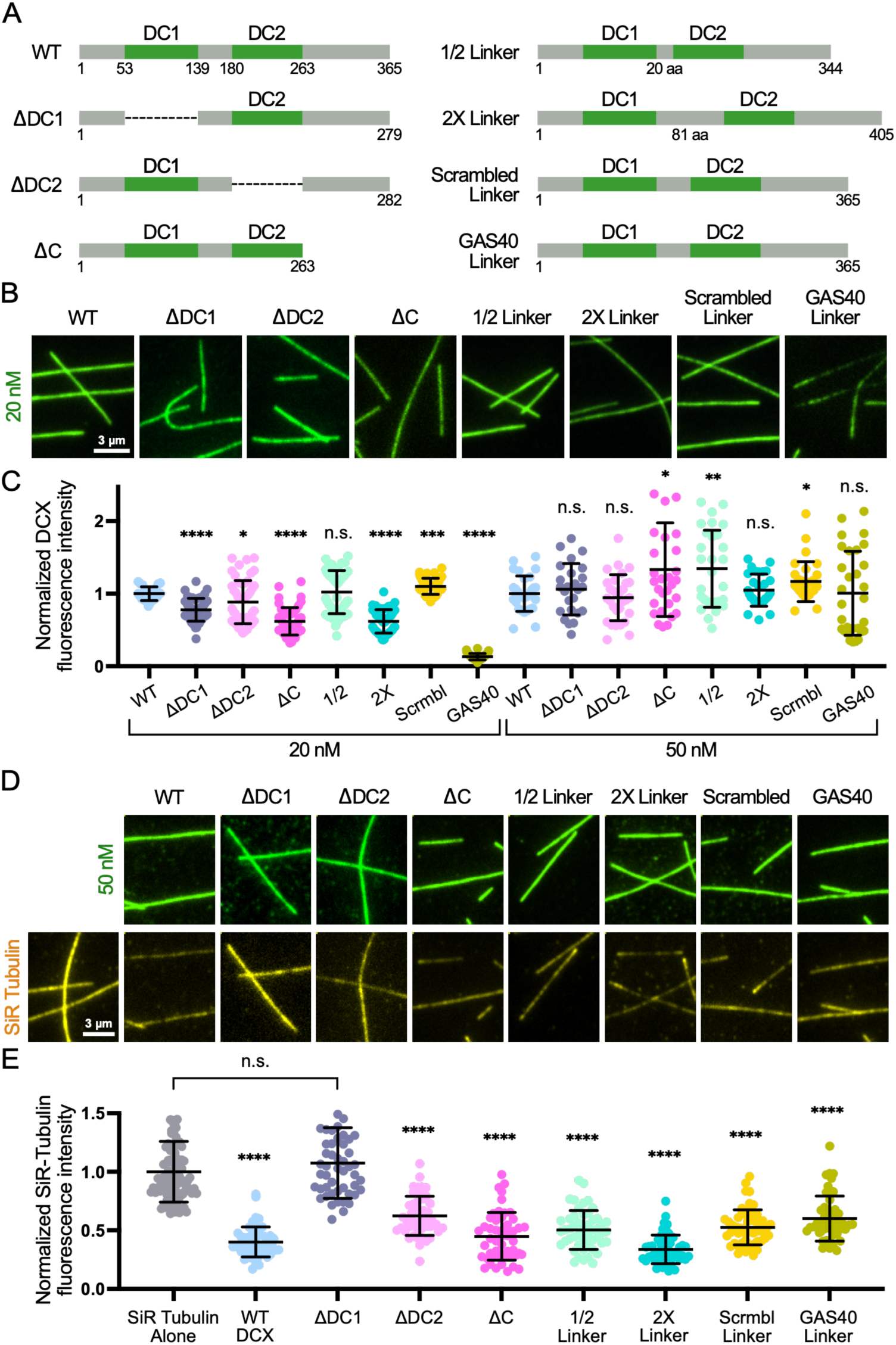
The DC1 Domain Is Necessary for DCX-Induced Lattice Compaction. **(A)** Diagrams of DCX constructs used for the domain analysis. **(B)** TIRF-M images of 20 nM of each DCX construct bound to taxol-stabilized microtubules. Scale bars = 3 µm; all images are the same size. **(C)** Quantification of normalized fluorescence intensity of each DCX construct on taxol-stabilized microtubules compared to the average fluorescence intensity of WT DCX at concentrations of 20 nM and 50 nM. Data are presented as mean ± s.d. For 20 nM: WT, 1.000 ± 0.096 (n = 25 microtubules); ΔDC1, 0.779 ± 0.158 (n = 50 microtubules); ΔDC2, 0.884 ± 0.297 (n = 50 microtubules); ΔC, 0.620 ± 0.188 (n = 48 microtubules); 1/2 Linker, 1.022 ± 0.296 (n = 50 microtubules); 2X Linker, 0.618 ± 0.161 (n = 50 microtubules); Scrambled Linker, 1.101 ± 0.111 (n = 50 microtubules); GAS-40 Linker, 0.132 ± 0.044 (n = 50 microtubules). **** indicates *p* < 0.0001. Other *p*-values compared to 20 nM WT: ΔDC2, 0.0147; 1/2 Linker, 0.6318; Scrambled Linker, 0.0002. For 50 nM: WT, 1.000 ± 0.243 (n = 30 microtubules); ΔDC1, 1.062 ± 0.354 (n = 25 microtubules); ΔDC2, 0.945 ± 0.317 (n = 30 microtubules); ΔC, 1.331 ± 0.646 (n = 30 microtubules); 1/2 Linker, 1.344 ± 0.530 (n = 29 microtubules); 2X Linker, 1.047 ± 0.221 (n = 30 microtubules); Scrambled Linker, 1.167 ± 0.274 (n = 30 microtubules); GAS-40 Linker, 1.006 ± 0.579 (n = 30 microtubules) from n = 2 independent experiments. *p*-values compared to 50 nM WT: ΔDC1, 0.4651; ΔDC2, 0.4554; ΔC, 0.0125; 1/2 Linker, 0.0028; 2× Linker, 0.4371; Scrambled Linker, 0.0156; GAS-40 Linker, 0.9581. **(D)** TIRF-M images of 1 µM SiR-tubulin (yellow) bound to taxol-stabilized microtubules in the absence (left) and presence of 50 nM of each DCX construct (green). Scale bars = 3 µm. **(E)** Quantification of SiR-Tubulin fluorescence intensity in the absence or presence of each DCX construct. Fluorescence intensity was normalized to SiR-tubulin alone. Data are presented as mean ± s.d. SiR Tubulin alone, 1.000 ± 0.259 (n = 100 microtubules); WT DCX, 0.401 ± 0.129 (n = 50 microtubules); ΔDC1, 1.075 ± 0.302 (n = 50 microtubules); ΔDC2, 0.623 ± 0.168 (n = 50 microtubules); ΔC, 0.449 ± 0.204 (n = 50 microtubules); 1/2 Linker, 0.503 ± 0.166 (n = 50 microtubules); 2X Linker, 0.337 ± 0.123 (n = 50 microtubules); Scrambled Linker, 0.526 ± 0.150 (n = 55 microtubules); GAS-40 Linker, 0.601 ± 0.192 (n = 50 microtubules) from n = 2 independent experiments. *p* = 0.1398 for SiR-Tubulin alone vs. ΔDC1. **** indicates *p* < 0.0001. All *p*-values were calculated using a two-tailed unpaired Student’s t-test. All graphs display all datapoints with lines indicating means ± s.d.

At 20 nM, all constructs exhibited decreased fluorescence intensity on taxol-stabilized microtubules compared to WT DCX, with the exception of the ½ Linker and Scrambled Linker constructs (Figure 3B-C). The GAS-40 Linker construct showed the most dramatic reduction in binding, with only 13.2% of WT intensity. At 50 nM, all constructs bound similarly to WT, suggesting that saturating densities were reached for all constructs at this concentration and indicating that the observed differences were due to altered binding affinities rather than loss of function or density of accessible sites on the microtubule (Figure 3C). To determine if these constructs could still compact the lattice, we used the SiR-tubulin displacement assay. Each construct was tested at 50 nM on taxol-stabilized microtubules, and SiR-tubulin intensity was measured and normalized to the SiR-tubulin alone control (Figure 3D-E). WT DCX reduced SiR-tubulin intensity by 60% compared to control levels, confirming lattice compaction. ΔDC1 failed to reduce SiR-tubulin intensity, indicating that ΔDC1 cannot compact the lattice. In contrast, all other constructs significantly reduced SiR-tubulin intensity compared to control. These results demonstrate that DC1 is important for lattice compaction, while DC2, the C-terminal region, and the linker domain are dispensable or redundant for this function.

### DC1 Enables DCX Self-Association on the Microtubule Lattice

Given that DC1 is necessary for lattice compaction but DC2 is not, we sought to understand the mechanistic differences between these two domains. We hypothesized that DC1-mediated lattice binding positions DC2 and the C-terminal region such that they can participate in intermolecular interactions with neighboring DCX molecules, facilitating self-association and lattice compaction. To test this hypothesis, we performed mixing experiments where 2.5 nM WT sfGFP-DCX was co-incubated with 100 pM of either WT mScarlet-DCX, mScarlet-DCX-ΔDC1, or mScarlet-DCX-ΔDC2 on taxol-stabilized microtubules (Figure 4A). WT mScarlet-DCX strongly co-enriched within WT sfGFP-DCX envelopes (Figure 4B-C). Similarly, mScarlet-DCX-ΔDC2 showed robust co-enrichment with WT envelopes. In stark contrast, mScarlet-DCX-ΔDC1 showed no enrichment within WT envelopes, and in fact exhibited a significantly higher fluorescence intensity outside of WT envelopes (Figure 4B-C). These results indicate that DC1 is important for DCX self-association on the microtubule lattice, as constructs lacking DC1 did not participate in cooperative envelope formation. Based on our findings, we propose a model (Figure 4D) in which DC1 anchors DCX to the microtubule lattice, allowing DC2 and the C-terminal to be available to interact with the DC2 and/or the C-terminal region of neighboring DCX molecules, facilitating self-association and lattice compaction that is potentially cooperative. In ΔDC2, DC1 can still bind the lattice, and the substantial intrinsically disordered region encompassing the linker and C-terminal segments (142 amino acids) remains available for interaction with neighboring WT DCX molecules. However, in ΔDC1, the shorter N-terminal region (N-terminal IDR + linker = 92 amino acids) and the C-terminal region are likely ill-positioned when DC2 is bound to the microtubule, and therefore less accessible for intermolecular interactions. This would explain why ΔDC1 cannot self-associate or compact the lattice, despite retaining DC2-mediated lattice binding. It is also consistent with prior NMR and cross-linking studies that reported transient interactions between these two domains^8,28^.

**Figure 4.**
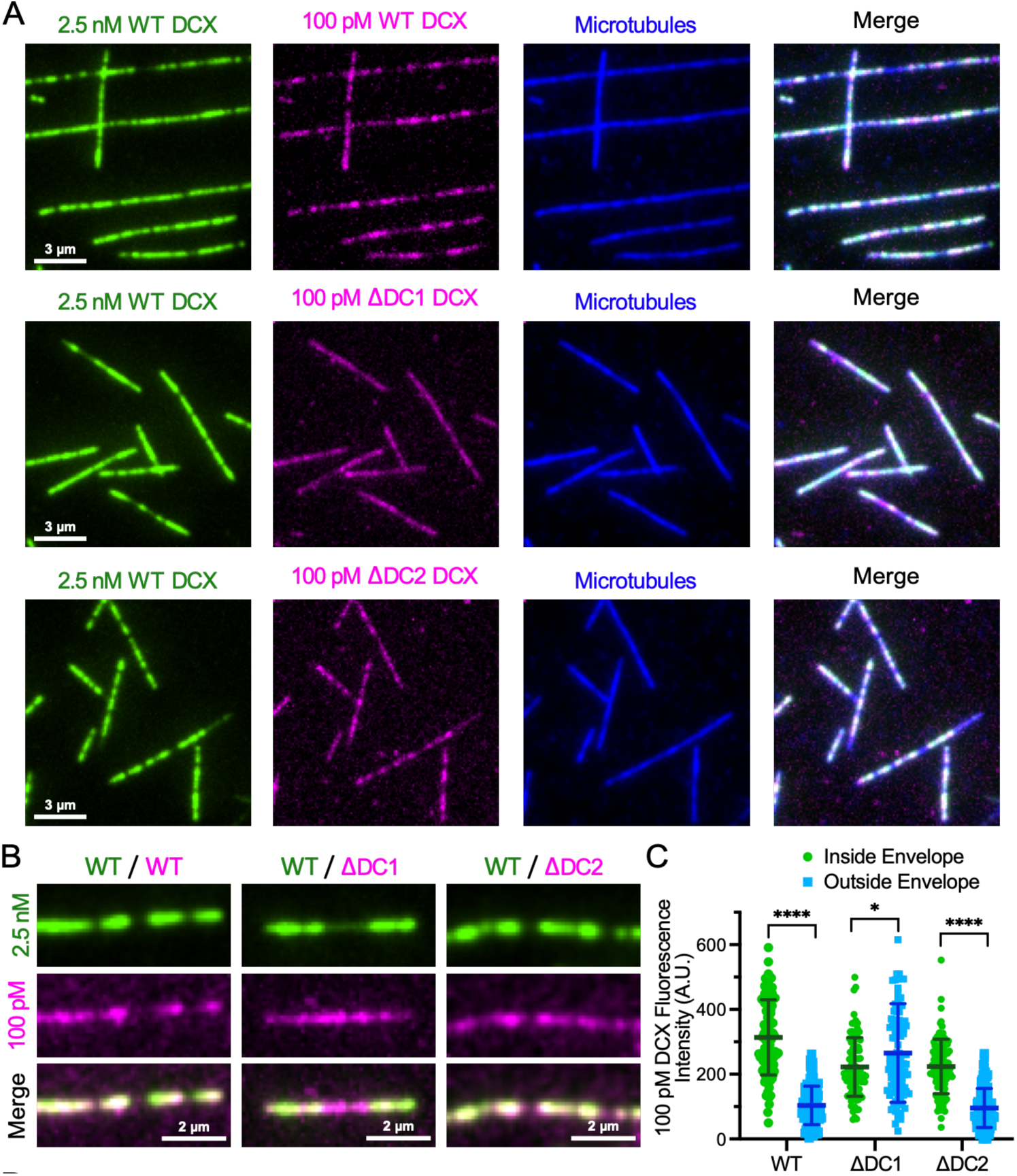
DC1 Is Required for DCX Self-Association on the Microtubule Lattice. **(A)** TIRF-M images of 2.5 nM WT sfGFP-DCX (green) with 100 pM WT mScarlet-DCX, 100 pM mScarlet-DCX-ΔDC1, or 100 pM mScarlet-DCX-ΔDC2 (magenta) on taxol-stabilized microtubules (blue). Scale bars = 3 µm. **(B)** Zoomed-in TIRF-M images showing miscibility between WT DCX and WT DCX, as well as between WT DCX and DCX-ΔDC2, but an anticorrelation between WT DCX and DCX-ΔDC1. Scale bars = 2 µm. **(C)** Quantification of fluorescence intensity of the 100 pM DCX construct inside vs. outside the 2.5 nM WT DCX envelope. Data are presented as mean ± s.d. For WT, inside the envelope, 313.7 ± 116.0; outside the envelope, 103.5 ± 59.5. For ΔDC1, inside the envelope, 222.1 ± 90.6; outside the envelope, 265.5 ± 152.2. For ΔDC2, inside the envelope, 223.8 ± 84.0; outside the envelope, 95.7 ± 60.5. n = 100 microtubules per condition from n = 2 independent experiments. *p* = 0.0153 for ΔDC1 inside vs. outside the envelope. **** indicates *p* < 0.0001. All *p*-values were calculated using a two-tailed unpaired Student’s t-test. The graph displays all datapoints with lines indicating means ± s.d.

### Structural Analysis Reveals Conformational Dynamics of DCX Domains

To gain molecular insights into the roles of DC1 and DC2 in DCX-induced lattice compaction, we performed atomistic molecular dynamics (MD) simulations on a microtubule fragment containing four intradimer and three interdimer interfaces, comparing APO (a bare, unoccupied microtubule fragment) to DC1- and DC2-bound states (Figure 5A). Principal component (PC) analysis revealed that the APO state predominantly exhibited lateral “breathing” motions, while DC1 and DC2 systems displayed a “shearing” motion at interdimer interfaces, along with a “hinge” motion at the interdimer interface (PC2 of DC1, PC1 of DC2), which are absent in APO^29^.

**Figure 5.**
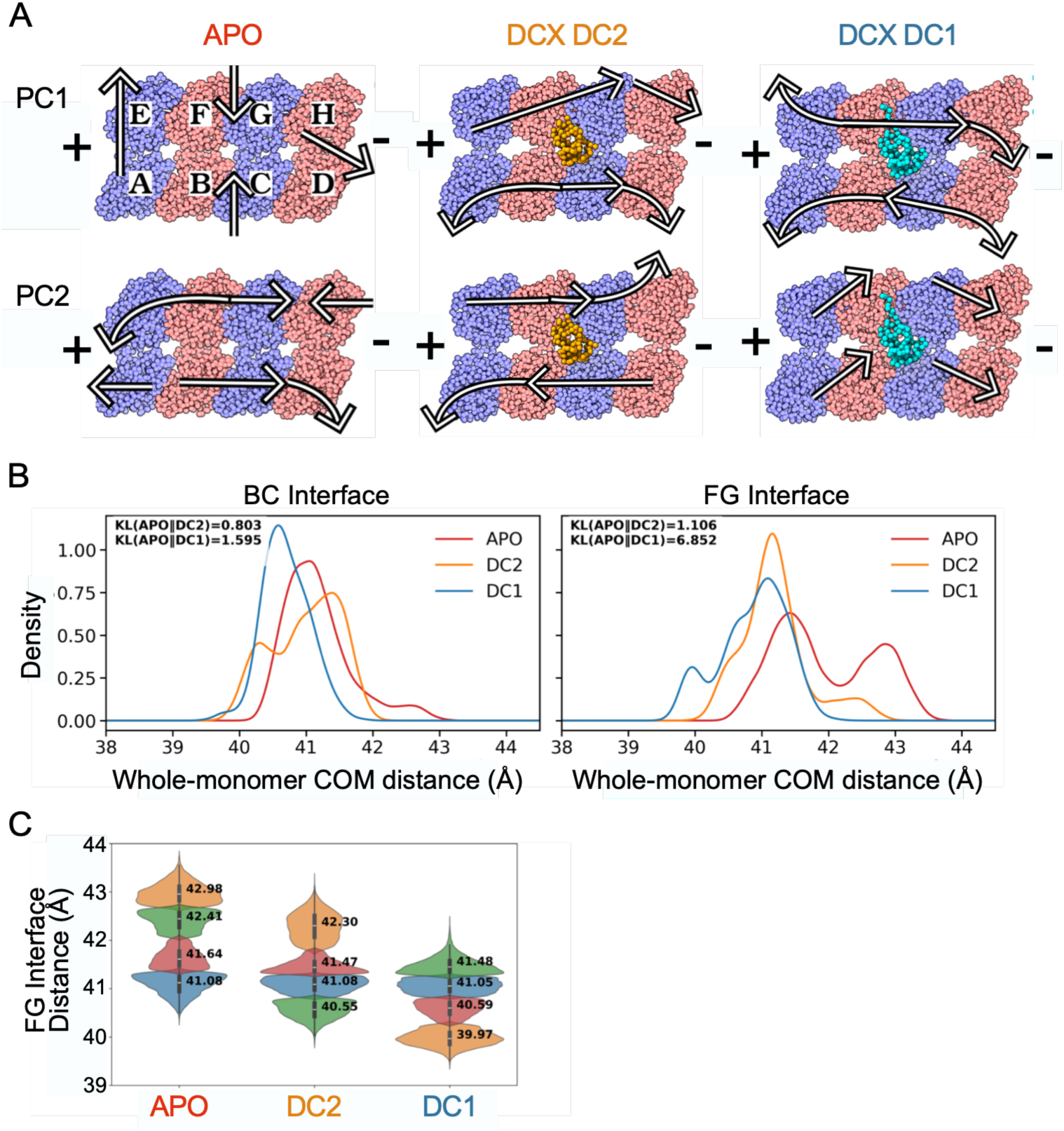
Structural analyses and center-of-mass distances between monomers for APO, DC1, and DC2 systems. **(A)** Molecular dynamics simulations of a two-protofilament microtubule fragment in the absence (APO) or presence of the DC1 or DC2 domain. Arrows depict the first two principal component motions, revealing collective movements of tubulin monomers including shearing (opposing protofilament sliding) and hinge motions at interdimer interfaces. **(B)** Probability distributions of center-of-mass (COM) distances between monomers at the BC and FG interdimer interfaces for APO (red), DC1 (blue), and DC2 (orange). Distributions shifted toward shorter distances indicate lattice compaction. Kullback–Leibler (KL) divergence scores quantify the extent to which each DCX-bound distribution differs from APO, with higher values reflecting greater conformational changes. **(C)** Violin plots of the COM FG interdimer distances for modes predicted by the Gaussian Mixture Model of each system’s distribution. The compacted states induced by DC1 represent new conformations not sampled by APO, with the mean of the most compacted DC1 state corresponding to ∼3 Å compaction relative to the most expanded APO state.

Quantification of center-of-mass (COM) distances between adjacent monomer pairs revealed that the most significant changes occurred at the interdimer interface (Figure 5B-C). Gaussian Mixture Model analysis showed that DC1 induced the most compacted state (39.97 Å) compared to APO (41.08 Å), representing ∼3 Å compaction at the most compacted state and ∼2 Å across metastable state mean spacings. DC2 also induced compaction (to 40.55 Å), but to a lesser extent than DC1. The hinge and shearing motions observed in DC1 restrict longitudinal tubulin movements, indicative of lattice stiffening.

We note that, compared to the experimental results, the MD simulations showed higher compaction values. This is likely due to simulating a small microtubule fragment. The fragment, which corresponds to the PDB entries for DCX domains, lacks normal lateral and longitudinal contacts with other protofilaments thus possibly allowing the DC domains to compact the lattice more than if the same subunits were part of a full microtubule. Importantly, modeled expanded states for APO and DC2 were comparable (∼43 Å), while DC1 still showed ∼1.5 Å compaction from the expanded states. At intradimer interfaces, minimal compaction was observed at the plus end, while the minus end showed modest compaction in both DCX-bound states at ∼ 0.75 Å (Figure S5A-B). Considering the interdimer compaction value, the simulations show that, on average, DCX domains compact the tubulin monomer-to-monomer spacing by ∼ 1.5 Å. These structural analyses support our biochemical findings that DC1 is the primary driver of lattice compaction through its ability to induce significant conformational changes at interdimer interfaces, while DC2 contributes to a lesser extent and primarily affects minus-end intradimer spacing.

### DCX Compaction Stiffens Microtubules and Inhibits Kinesin-1 and αTAT1 Binding

We next sought to investigate the functional consequences of this compaction activity. Recent *in vivo* studies have shown that DCX proteins reinforce microtubules by increasing their rigidity for specific cellular contexts^15,30^. To test the effect on microtubule mechanical properties, we used an assay where taxol-stabilized microtubules with reduced levels of biotinylated tubulin were sparsely anchored to a streptavidin-coated surface, causing them to adopt curved conformations. Using a FRAP laser, we severed the middle of microtubules coated with either DCX or MAP4 (Figure 6A). Successful severing was confirmed by the immediate loss of fluorescence signal at the cut site, distinguishing cutting from photobleaching. Images were acquired before severing and at 2 s and 15 s post-severing. Upon severing, DCX-coated microtubules immediately straightened and remained straight for the remainder of the movie, while MAP4-coated microtubules remained in their curved state. By calculating the curvature index at 15 s post-cut, (L₂-L₁)/L₂, where L₁ is the end-to-end distance and L₂ is the contour length, we found that MAP4-coated microtubules exhibited significantly higher curvature than DCX-coated microtubules (Figure 6B). These results are indicative of increased microtubule stiffness upon DCX binding.

**Figure 6.**
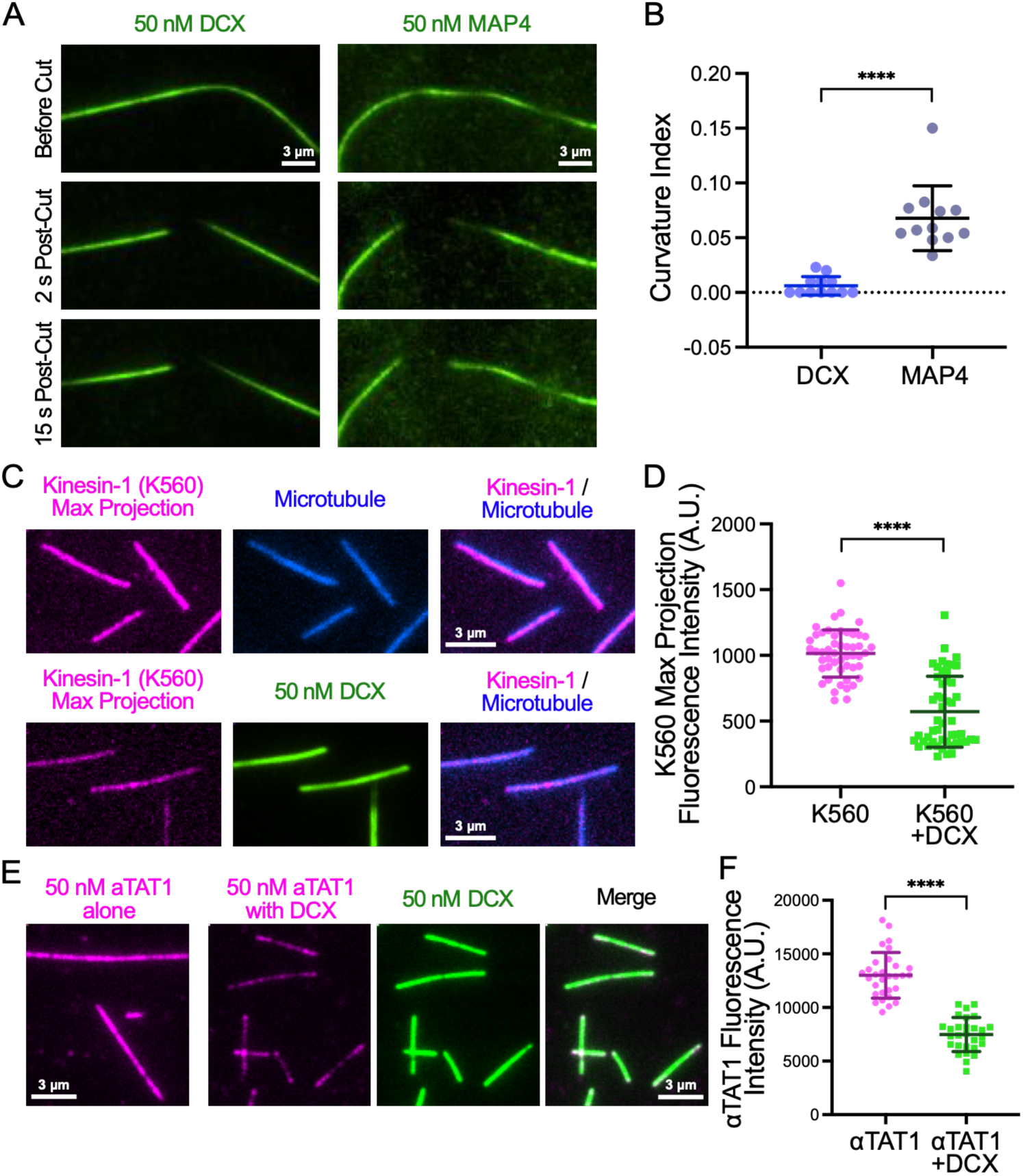
DCX Compaction Stiffens Microtubules and Inhibits Kinesin-1 Motility. **(A)** TIRF-M images showing a time course of DCX-coated and MAP4-coated taxol-stabilized microtubules before FRAP laser severing and 2 s and 15 s after cutting. Following laser cutting, DCX-coated microtubules rapidly straighten, whereas MAP4-coated microtubules remain curved and flexible. Scale bars = 3 µm. **(B)** Quantification of microtubule curvature post-cut. Curvature index was calculated as [(*L*_2_ − *L*_1_)/*L*_2_], where *L*_1_is the end-to-end distance and *L*_2_is the contour length of the microtubule. Higher values indicate greater curvature. Means ± s.d.: 0.006 ± 0.008 for DCX and 0.068 ± 0.030 for MAP4, from n = 12 microtubules per condition. **(C)** TIRF-M max projection images of 5 nM KIF5B (kinesin-1) (pink) with 1 mM ATP in the absence (top row) or presence (bottom row) of 50 nM sfGFP-DCX (green) on taxol-stabilized microtubules (blue). Max projections were generated from 3 min movies. Scale bars = 3 µm. **(D)** Quantification of K560 max projection fluorescence intensity in the absence and presence of sfGFP-DCX on taxol-stabilized microtubules. Means ± s.d.: 1014.2 ± 179.2 in the absence of DCX, 571.6 ± 270.2 in the presence of DCX, from n = 50 microtubules per condition from n = 2 independent experiments. **(E)** TIRF-M images of 50 nM αTAT1 (pink) in the absence (left) or presence (right) of 50 nM sfGFP-DCX (green) on taxol-stabilized microtubules. Scale bars = 3 µm. **(F)** Quantification of αTAT1 fluorescence intensity in the absence and presence of sfGFP-DCX on taxol-stabilized microtubules. Means ± s.d.: 13005 ± 2132 in the absence of DCX, 7471 ± 1587 in the presence of DCX, from n = 30 and 28 microtubules, respectively, from n = 2 independent experiments.

Given that DCX compacts the lattice and kinesin-1 is known to prefer an expanded lattice state^31,32^, we examined whether DCX allosterically inhibits kinesin-1 motility. We imaged 5 nM KIF5B-K560 (kinesin-1, aa 1-560) in the absence or presence of 50 nM DCX and found that in the presence of DCX, the presence of K560 on microtubules was substantially reduced (Figure 6C-D). This inhibition is likely allosteric, as the DCX and kinesin-1 binding sites on the microtubule do not overlap^33^. Instead, DCX-mediated lattice compaction apparently creates a conformational state that is incompatible with kinesin-1 binding and motility^18^.

We previously demonstrated that α-tubulin acetyltransferase 1 (αTAT1) preferentially binds and acts on expanded microtubule conformations^24^. We thus investigated whether DCX-induced compaction affects αTAT1 binding. DCX reduced αTAT1 binding to microtubules by approximately 43% (Figure 6E-F), indicating that DCX antagonizes αTAT1 binding.

Together, these results establish a mechanistic framework in which DCX preferentially recognizes and binds to compacted microtubule lattices, but can also bind expanded lattices and compact them through cooperative self-association into protein envelopes. This ability is highly similar to that of the orthogonal neuronal MAPs tau and MAP2^22^, despite no sequence or structure conservation between these MAP families. This compaction both stiffens the microtubule and allosterically regulates kinesin-1 and aTAT1 access by creating a lattice state unfavorable for binding.

## DISCUSSION

In this study, we have uncovered a mechanistic foundation for DCX’s diverse cellular functions, from neuronal migration to growth cone advance in soft environments. DCX preferentially binds compacted GDP-lattices, but it can also bind expanded lattices and compact them through a DC1-dependent mechanism that involves shearing and hinge motions at interdimer interfaces. Compaction is associated with stiffening of taxol-stabilized microtubules and inhibition of kinesin-1 and αTAT1 binding.

Our finding that DC1 is necessary to position DCX for efficient lattice compaction and self-association, while DC2 is dispensable, raises intriguing questions about functional diversification within the DCX superfamily. Most human doublecortin family members contain two DCX domains, yet DCDC1 and DCLK3 possess only single DCX domains that show greatest similarity to DC2^34^. Based on our results, we propose that this single-domain architecture may have evolved to specifically select DC2-mediated lattice binding that avoids cooperative self-association and lattice compaction. The ability of DC1-containing DCX isoforms to compact microtubules is thus a selective function that depends on on physiological demands. Our findings extend prior observations that DCX preferentially binds certain microtubule lattice conformations^11,12^ and that DCX competes with taxol for control of lattice spacing^26^. Our study provides the first direct demonstration of DCX-induced lattice compaction using three orthogonal methods and identifies DC1 as the necessary domain for this activity.

It is striking that DCX, tau, and MAP2, which have entirely different domain architectures and binding sites along the microtubule, have seemingly converged on a common mechanism of cooperative binding and lattice compaction. Tau and MAP2 bind through intrinsically disordered repeat domains contacting the outer surface of protofilaments^35,36^, while DCX binds between protofilaments at interdimer interfaces^6^. Despite these fundamental differences, both protein families form cooperative protein envelopes that induce similar lattice compaction^11,21,22^, and are equally sensitive to Zn^2+^ ^37^. This remarkable convergence suggests that compacting microtubules and regulating motor access is a fundamentally important function that has evolved independently in distinct MAP families.

We previously reported that DCX and kinesin motors compete for microtubule access^18^. Our current results suggest an allosteric mechanism beyond steric competition. By inducing lattice compaction, DCX creates a conformational state unfavorable for kinesin-1 binding, which is supported by observations that kinesin-1 expands the lattice^31,32,38^. Interestingly, the DCLK1 homolog, ZYG-8, has been shown to inhibit kinesin-5 that promotes spindle stability in *C. elegans* oocytes^39^, indicating that the DCX superfamily may regulate multiple kinesins depending on cellular context. DCX also negatively regulates dynein-mediated retrograde transport^20^, suggesting that DCX may coordinate regulation of both anterograde and retrograde motors.

The observation that DCX compaction may stiffen microtubules provides a mechanistic explanation for prior observations that DCX reinforces the microtubule cytoskeleton and promotes growth cone advance in soft environments^15^. By stiffening microtubules or maintaining the stiffness of endogenous microtubules, DCX may reinforce the growth cone cytoskeleton to counteract actomyosin-generated contractile forces, fostering neuronal migration and growth cone advance. This is consistent with work in *C. elegans* showing that ZYG-8 increases microtubule rigidity and that reduced rigidity leads to impaired spindle positioning^30^. Our finding that DCX compaction may serve to stiffen microtubules provides a potential molecular mechanism for these observations and suggests this function is evolutionarily conserved.

In our molecular dynamics simulations, we found that DC1 binding induces ∼2 Å average compaction at the interdimer interface compared to APO, while DC2 has a more modest effect. This compaction results from shearing motion, collective sliding of monomers from one protofilament away from its lateral neighbor, and hinge motion, where end dimers move together forming a hinge at the interdimer interfaces (BC and FG). Notably, compacted states at interdimer interfaces induced by DCX represent new conformations not explored by APO fluctuations, while compacted states at intradimer interfaces are already sampled in APO. This distinction highlights that the primary effect of DCX binding is stabilizing specific compacted conformations at interdimer interfaces, consistent with the DCX binding site between protofilaments^6^. The broader APO distributions suggest the lattice intrinsically samples wider conformations, and DCX induces stabilized compacted states, reducing lattice heterogeneity. The hinge and shearing motions observed with DC1 restrict longitudinal tubulin movements, which is predicted to increase lattice rigidity and provides a structural basis for the experimentally observed increased stiffness.

The ability of DCX to compact the microtubule lattice has important implications for the regulation of tubulin post-translational modifications (PTMs)^40^. Recent work has shown that the detyrosination machinery, VASH/SVBP and MATCAP1, preferentially acts on expanded microtubule conformations^41,42^. Likewise, MAP7, a microtubule expander, promotes acetylation by αTAT1^24^. Additionally, DCX restricts neuronal branching by regulating α-tubulin polyglutamylation^19^, suggesting that DCX coordinates lattice compaction with post-translational modification (PTM) pathways. Given that the basic microtubule lattice exists in a dynamic equilibrium between expanded and compacted states, DCX-induced compaction may fundamentally alter the accessibility of modifying enzymes to the microtubule surface. By stabilizing the compacted conformation, DCX could potentially bias the lattice away from detyrosination and acetylation, which favor expanded conformations, while promoting polyglutamylation. These observations collectively build a framework in which MAPs regulate the PTM landscape of the microtubule cytoskeleton through the control of lattice conformation. We further propose that MAP envelope formation could locally alter lattice conformation, providing a mechanism for spatial regulation of microtubule PTMs.

In summary, we have established that DCX compacts the microtubule lattice through a DC1-dependent mechanism involving cooperative self-association and conformational changes at interdimer interfaces, which stiffens microtubules and allosterically inhibits kinesin-1 motility. The convergence of these functions in evolutionarily distinct MAP families highlights their fundamental importance for cellular mechanics and motor regulation. Our findings provide a conceptual basis for understanding how DCX contributes to neuronal development and disease through the control of microtubule lattice conformation.

## ACKNOWLEDGEMENTS

The authors wish to thank all the members of the Ori-McKenney and McKenney laboratories for their kind help and feedback. This work is supported by the NIH grants R35GM133688 to K.M.O.-M., R35GM124889 to R.J.M., R35GM118139 to Y.E.G., NSF grant MCB-2527485 to R.D. and K.M.O.-M., EMBO Postdoctoral Fellowship (ALTF 1069-2024) to V.S., the Czech Science Foundation (GACR) grant 25-17813S to M.B., and Z.L. is supported by the grant ERC-2022-SYG 101071583 from the European Research Council. The optical trapping system used in this study was supported by NIH Grant 1S10OD036332. We acknowledge institutional support from CAS (RVO: 86652036), Imaging Methods Core Facility at BIOCEV supported by the MEYS CR [LM2023050, Czech-BioImaging] and European Regional Development Fund-Project”UP CIISB” [No. CZ.02.1.01/0.0/0.0/18_046/0015974].

## AUTHOR CONTRIBUTIONS

R.A., R.J.M., and K.M.O.-M. conceived the project and designed the TIRF-M experiments. R.A. purified the proteins, performed the *in vitro* TIRF-M experiments, and R.A. and M.F.H. analyzed the data. L.B.S. performed the αTAT1 in vitro experiments and analyzed the data. J.S., C.S.C., and R.D. designed the molecular dynamics simulations. J.S. and C.S.C. performed the molecular dynamics simulations and analyzed the data. H.S. and Y.E.G. designed the optical trapping experiments, and H.S. performed the experiments and analyzed the data. M.B., Z.L., and V.S. designed the speckle tubulin assays, and V.S. performed the experiments and analyzed the data. S.E.L. performed the cryo-EM analysis. R.J.M. performed the laser severing assays, and Y.S. analyzed the data. R.A. and K.M.O.-M. wrote the manuscript with input from all authors. All authors reviewed and edited the manuscript.

## MATERIALS AND METHODS

### Protein Constructs and Purification

For in vitro experiments, DCX constructs were cloned into pET28a vectors using Gibson assembly. All DCX constructs contain a N-terminal cassette consisting of a superfolder GFP (sfGFP) fluorophore and StrepII tag. In addition, we made wild type DCX, DCX-ΔDC1, and DCX-ΔDC2 constructs tagged N-terminally with an mScarlet fluorophore and a StrepII tag. For preparation of Kinesin-1 (KIF5B K560), a DNA fragment encoding human KIF5B aa 1-560 was codon-optimized for E. coli and synthesized by gBlocks (Integrated DNA Technologies, Coralville, IA, USA). Gibson assembly was again used to clone the fragment into a pET28a vector with a C-terminal cassette consisting of an mScarlet fluorophore and StrepII tag. K560 was then expressed and purified as previously described^43^. αTAT1 (human isoform 4 modified from Addgene plasmid 27099) was cloned into a pET28a vector with a C-terminal mScarlet-strepII cassette.

For bacterial expression and purification of all DCX variants and αTAT1, BL21 cells were grown at 37°C until an OD of 0.6 was reached and protein expression was subsequently induced with 0.1 mM isopropyl-beta-D-thiogalactopyranoside (IPTG). Cells were grown overnight at 18°C, pelleted at 3700g for 15 min prior to flash freezing in liquid nitrogen and storing at -80°C. Cell pellets were resuspended in lysis buffer (50 mM Tris pH 8.0, 150 mM K acetate, 2 mM Mg acetate, 1 mM ethylene glycol tetraacetic acid (EGTA), 10% glycerol) and supplemented with protease inhibitor cocktail (Roche), 1 mM dithiothreitol (DTT), 1 mM phenylmethylsulfonyl fluoride (PMSF), and DNAseI. Cells were homogenized via a dounce homogenizer and passed through an Emulsiflex press prior to addition of 0.1% Triton-X 100 and cleared via centrifugation at 22,700g for 20 min at 4°C. Clarified lysate from bacterial expression was passed over a column with Streptactin-XT Superflow resin (Qiagen). After incubation for 1 hr at 4°C in the resin, the column was washed with lysis buffer supplemented with 1 mM DTT and 1 mM PMSF. Protein was then eluted with lysis buffer supplemented with 1 mM DTT, 1 mM PMSF and 100 mM d-Biotin (CHEM-IMPEX) pH 8. Eluted proteins were further purified via anion (HiTrap Q) or cation (HiTrap SP HP) exchange chromatography in lysis buffer as appropriate or via size exclusion chromatography with a Superdex 200 10/300 GL (Cytiva) column equilibrated in lysis buffer using a Bio-Rad NGC system. The resulting preparations were analyzed by sodium dodecyl sulfate-polyacrylamide gel electrophoresis (SDS-PAGE) and peak fractions were pooled and concentrated via Amicon concentrators, flash frozen in liquid nitrogen and stored at -80°C. Protein concentration was measured by Nanodrop One (ThermoFisher) based on absorbance of attached fluorescent tag.

### Microtubule Assembly

Tubulin was isolated from porcine brain using the high-molarity PIPES procedure as previously described^44^.

#### Taxol-lattice microtubules

For TIRF-M experiments, taxol-stabilized microtubules were polymerized utilizing a mixture of native tubulin, biotin-tubulin, and fluorescent-tubulin purified from porcine brain (∼10:1:1 ratio) in BRB80 buffer (80 mM PIPES, 1 mM MgCl2, 1 mM EGTA, pH 6.8 with KOH) by incubating with 1 mM GTP for 30 min at 37°C. Polymerized MTs were then stabilized with 20 μM taxol and incubated for another 30 min before pelleting over a 25% sucrose cushion at 15,000g to remove unpolymerized tubulin. After centrifugation, the pellet was carefully resuspended and kept in BRB80 with 10 μM taxol. For speckle experiments, taxol-lattice microtubules were polymerized from 4 mg/ml tubulin (2% biotin-labeled tubulin and 0.133% Atto647-labeled tubulin) as described above. For TIRF-M imaging experiments, taxol-stabilized microtubules were diluted into BC Buffer (BRB80 supplemented with 1 mg ml−1 bovine serum albumin (BSA), 1 mg ml−1 casein) and 10 μM taxol prior to imaging.

#### GMPCPP-lattice microtubules

GMPCPP-lattice microtubules were polymerized utilizing a mixture of native tubulin, biotin-tubulin, and fluorescent-tubulin purified from porcine brain (∼10:1:1 ratio) in BRB80 supplemented with 1 mM GMPCPP (Jena Bioscience, NU-405) and 1 mM DTT and incubating for 20 min at 37°C. The polymerized GMPCPP microtubules were pelleted over a 25% sucrose cushion at 15,000g to remove unpolymerized tubulin. After centrifugation, the pellet was carefully resuspended and kept in BRB80 supplemented with 1 mM DTT. For TIRF-M imaging experiments, GMPCPP microtubules were diluted in BRB80 prior to imaging.

#### GDP-lattice microtubules polymerized from GMPCPP seeds

GMPCPP seeds were prepared as described above using a mixture of biotin-tubulin and unlabeled tubulin. Seeds were diluted and allowed to adhere to the coverslip surface, then 20 μM unlabeled tubulin in BRB80 with 1 mM GTP and 1 mM DTT was flowed into the chamber and incubated for 10 min at 37°C to allow elongation from the seeds.

### TIRF Microscopy

Movies and images were acquired utilizing a NIS-Elements software-controlled (AR, Version 5.20.2) Nikon Eclipse Ti2 Microscope (1.49 numerical aperture, ×100 objective) equipped with a TIRF illuminator and Andor iXon charge-coupled device electron multiplying camera. For speckle experiments, total internal reflection fluorescent (TIRF) microscopy assays were performed on an inverted widefield microscope (Nikon Eclipse Ti2), equipped with motorized XY-stage, Perfect Focus System, Nikon Apo TIRF 60x oil immersion objective (NA 1.49), 1.5x tube lens, and PRIME BSI camera (Teledyne Photometrics). Microtubules were visualized using interference reflection microscopy (IRM). Fluorescent microtubule speckles and DCX were visualized using a EM610/75 and EM435-485 filter turret, respectively.

#### Buffers and Conditions

Flow chambers containing immobilized microtubules were assembled utilizing acid-washed coverslips as described previously^45^. For speckle experiments, TIRF chambers were assembled from glass coverslips silanized with Hexamethyldisilazane (HMDS, Sigma, #379212) and thin strips of parafilm melted in between two coverslips. The chambers were incubated with 20 μg/ml anti-biotin antibodies (#B3640, Sigma) for 5 min, followed by 1% F-127 (#P2443, Sigma, 1% in PBS) for at least 30 min. All experiments were performed in assay buffer (60 mM HEPES pH 7.4, 50 mM K-acetate, 2 mM Mg-acetate, 1 mM EGTA, and 10% glycerol) supplemented with 0.1 mg/mL biotin-BSA, 0.5% Pluronic F-168, and 0.2 mg/mL κ-casein (Sigma), unless otherwise stated. 10 μM taxol was added to assay buffer when utilizing taxol-stabilized microtubules. All experiments were conducted at room temperature, unless otherwise stated.

#### DCX binding to different microtubule lattices

Taxol-stabilized or GMPCPP microtubules were immobilized on the coverslip surface and unbound microtubules were removed with assay buffer. sfGFP-DCX was diluted to the final concentrations indicated in the main text in assay buffer supplemented with 2 mM Trolox, 2 mM protocatechuic acid and 50 nM protocatechuate-3,4-dioxygenase and added to the TIRF chamber. For GDP-lattice experiments, GMPCPP seeds were immobilized and GDP-lattice microtubules were polymerized as described above, then sfGFP-DCX was added as described.

#### SiR-Tubulin Assay

SiR-Tubulin (Cytoskeleton, Inc.) was resuspended in DMSO as per manufacturer instructions. For TIRF-M, taxol-stabilized or GMPCPP microtubules were prepared as described above, diluted and immobilized in TIRF chambers. SiR-Tubulin and appropriate DCX construct were then diluted together in assay buffer supplemented with 2 mM Trolox, 2 mM protocatechuic acid and 50 nM protocatechuate-3,4-dioxygenase and added to the TIRF chamber to the final assay concentration stated in the main text before imaging on TIRF-M within five minutes.

#### Kinesin-1 on microtubules

GMPCPP- or taxol-lattice microtubules were prepared as above and immobilized on the coverslip surface. Unbound microtubules were removed with assay buffer and TIRF-M objective was set at 30°C. mScarlet-Kinesin 1_K560 and WT DCX-sfGFP were diluted to the final concentrations listed in the main text in BRB80 supplemented with 0.1 mg/mL biotin-BSA, 0.5% Pluronic F-168, 0.2 mg/mL κ-casein (Sigma). 10 μM taxol was added to the assay buffer if taxol-stabilized microtubules were being used. 1 mM ATP and 2 mM Trolox, 2 mM protocatechuic acid and 50 nM protocatechuate-3,4-dioxygenase were also added to the reaction prior to imaging. Imaging of mScarlet-Kinesin1_K560 was conducted at 2.5 Hz for 3 min.

#### Envelope Enrichment Assay

For envelope enrichment assays, taxol-stabilized microtubules were prepared as above and immobilized on the coverslip. Unbound microtubules were removed with assay buffer. WT DCX-sfGFP and mScarlet-WT DCX, mScarlet-DCXΔDC2, or mScarlet-DCXΔDC1 were diluted and mixed in assay buffer supplemented with 2 mM Trolox, 2 mM protocatechuic acid and 50 nM protocatechuate-3,4-dioxygenase to final concentrations listed in the main text prior to flowing into TIRF chamber and imaging. Imaging of WT DCX-sfGFP and mScarlet-DCX constructs was conducted at 10 Hz for one minute. Envelope boundaries were defined as contiguous regions of microtubule where sfGFP-DCX fluorescence intensity exceeded a threshold set at 50% of the maximum intensity for that microtubule, and “outside” regions were adjacent segments of the same microtubule where intensity remained below this threshold.

#### Microtubule Lattice Compaction by Speckle Tracking

Speckled taxol-lattice microtubules were prepared as described above and immobilized in TIRF chambers. Before imaging, chambers were equilibrated in TIRF assay buffer AB (50 mM HEPES pH 7.4, 1 mM EGTA, 2 mM MgCl2, 75 mM KCl, 10 mM dithiothreitol, 0.02 mg/ml casein, 10 μM taxol, 1 mM Mg– ATP, 20 mM D-glucose, 0.22 mg/ml glucose oxidase and 20 μg/ml catalase). Next, DCX diluted in AB to the desired concentration was added to the chamber and imaged using TIRF microscopy. Microtubule lattice compaction by DCX was analyzed as previously described^22^. In short, atto647-labelled microtubule speckles were tracked using FIESTA tracking software (v1.6.0) to determine the distance between two neighboring speckles. Speckle spacing was averaged over five frames before DCX addition and compared to the average spacing of the same speckles after 5 min of DCX incubation (5 frames). Fluorescence images of DCX were then used to correlate the speckle localization to the position of the DCX envelopes on the microtubule lattice. For each microtubule, compaction values were averaged separately for speckles located within DCX envelopes or outside DCX envelopes.

### Optical Tweezers

Correlative force measurements and microscopy were performed on an optical tweezers setup equipped with confocal fluorescence imaging and microfluidic system (c-Trap, LUMICKS B.V.). The microfluidic system was passivated using casein (0.1 % in PBS buffer, pH 7.5). The trap stiffness (0.4 pN/nm) was calibrated using force calibration in Bluelake (v.2.9, LUMICKS B.V.) under the zero-flow condition. The experiments were performed at room temperature. Data was acquired and exported using the Bluelake software (v.2.9) and analyzed using Lakeview (v. 1.3.2). Forces were measured along the microtubule *via* the Bluelake calibration. Inter-bead distances, representing the length of the tethered microtubule, were determined by calculating the edge-to-edge distance (center-to-center distance less 1× the diameter of the beads).

#### Microtubule lattice compaction

Two streptavidin silica beads (1.76 μm, Lumicks) were captured in two separate traps. The beads were then moved into the channel containing Alexa647-labeled microtubules (in SPR-90 buffer supplemented with 2 μM taxol). Following microtubule binding to one bead, the assay was moved into buffer only channel, where the microtubule was tethered between two beads using flow thereby forming a “dumbbell”. The microtubule, initially in a relaxed state (0 pN), was then stretched by moving one of the beads at a pulling velocity of 0.1 μm/s until the applied force reached 25 pN. Each microtubule was subjected to a minimum of four stretch-relax cycles using the built-in force spectroscopy loop in Bluelake software, while its length was continuously monitored. Continuous 2D confocal scans were acquired to monitor the Alexa647-labeled microtubule under 638 nm excitation. The construct was then moved into the channel containing 50 nM sfGFP-DCX or sfGFP-MAP4 and the change in microtubule length was subsequently monitored for an additional 100 s under the same experimental conditions following the stretch-relax cycles. Continuous 2D confocal scans were acquired under 488 nm excitation to monitor the binding of DCX or MAP4 to the microtubule indicated by an increase in sfGFP fluorescence on the microtubule. Microtubule lattice compaction was quantified as the change in microtubule length measured at an applied force of 25 pN (Figure 2A-D). The percent compaction was calculated from the change in inter-bead distance upon protein binding relative to the initial length in buffer (Figure 2E). Representative time traces showed consistent photon counts in the red channel confirming the presence of the Alexa647-labeled microtubule throughout the experiment, while the increase in blue-channel photon counts indicated binding of GFP-tagged DCX or MAP4 to the microtubule.

### Cryo-Electron Microscopy

For cryo-EM experiments, 20 μM unlabeled porcine brain tubulin (Cytoskeleton #T240) was polymerized at 37°C for 2 h in BRB80 (80 mM PIPES pH 6.8, 1 mM MgCl2, 1 mM EGTA) supplemented with 3 mM GTP. An equal volume of BRB80 supplemented with 10 μM taxol was added, and the polymerized microtubules were left at room temperature overnight. Directly before grid preparation, the microtubules were pelleted at 20,000 r.c.f. for 10 min and resuspended to 3 μM in fresh BRB80.

Microtubules and DCX were pre-incubated before being added to electron microscopy grids. A mixture of 3 μL 3 μM microtubules and 1 μL 8 μM DCX was incubated at room temperature for 20 min. For the microtubule-only dataset, DCX was replaced with BRB80. During incubation, R1.2/1.3 Au300 Quantifoil EM grids were glow-discharged for 40 s. Then, 4 μl microtubule-DCX mixture was applied to the grid in a Vitrobot Mark II (TFS) set to 100% humidity and 22°C. After 30 s, the grid was blotted for 4.5 s and plunged into liquid ethane. The grids were loaded into a Polara cryo-electron microscope (TFS) operating at 300 kV. Images were collected semi-automatically in SerialEM with a pixel size of 1.35 Å px−1, defocus ranging from −2 to −3 μm, and a dose rate of 25 e−1 px−1 s−1 for a 1.5-s exposure. Roughly forty images were taken for each dataset. Movie stacks were aligned in MotionCorr2. Straight and uninterrupted microtubule segments longer than 2.5 μm were then manually picked in RELION 3.0, with the coordinates used to crop out each microtubule. In FIJI, each microtubule segment was then rotated to line up with the y-axis of the image and the Fast Fourier Transform (FFT) was calculated. The spatial frequency of the peak in intensity of the 4 nm (J.) reflection, corresponding to the longitudinal lattice spacing, was measured manually in FIJI (Figure 2F).

### Laser Severing and Stiffening Assay

For laser severing experiments, taxol-stabilized microtubules were prepared and sparsely anchored to the coverslip surface to allow curved conformations. Microtubules were coated with either sfGFP-DCX or sfGFP-MAP4 at saturating concentrations (50 nM). A FRAP laser (405 nm) was used to sever the middle of individual microtubules. Images were acquired before severing and every second afterwards for 30 seconds (Figure 6A). Curvature index was calculated as [(L₂-L₁)/L₂], where L₁ is the end-to-end distance and L₂ is the contour length of the microtubule, measured using ImageJ (Figure 6B).

### Molecular Dynamics Simulations

The two cryo-EM structures of human beta-3 tubulin (TUBB3) with GDP and human alpha-1b tubulin (TUBA1B) with GTP and Mg2+ were obtained to construct a two protofilament microtubule complex with two tubulin dimers in each protofilament^46,47^. In PyMol, the human DC1 (6REV) and DC2 (6RF2) domains bound between four Bos taurus tubulin dimers of a 13-protofilament GDP-microtubule were used to align the human tubulin to the binding regions of DC1 and DC2^6,48^. We used the Modeller program in ChimeraX was used to better align the tubulin structures and avoid steric clashes due to alignment^49^. Molecular dynamics (MD) simulations using the AMBER99DB-ILDN force field were run on three different systems using GROMACS 2024: a two protofilament system with no DCX domains bound (APO state), with the DC1 domain bound (DC1 state), and with the DC2 domain bound (DC2 state)^50^. Each system (∼230,000 atoms) was placed in a rectangular box with the TIP3P water model as the solvent and neutralized with sodium ions^51^. Restraints on the edges of the four tubulin were added at the microtubule lattice ends to mimic additional bound tubulin and to prevent unwanted bending. For each system, five separate 500 ns trajectories were run and root mean square deviation (RMSD) of the protein backbone was calculated to evaluate convergence.

Principal component analysis (PCA) was performed on all systems to determine how doublecortin influences microtubule dynamics. The equilibrated parts of the trajectories in a system were concatenated, and the dimensions were reduced to capture the collective motions with the most variance. The covariance matrix was calculated using the carbon alpha atoms with GROMACS (gmx covar), and the first two principal components (gmx anaeig) were analyzed for each system (Figure 5A).

The center of mass (COM) distance between monomers at each interface was calculated using the MDanalysis library^52^. SciPy was used to make kernel density estimate probability distributions for each microtubule state (APO, DC2, DC1)^53^. The Kullback-Leibler divergence between each distribution in each chain pair was calculated^54^, where a KL value at 0 indicates identical distributions and higher KL values indicate distributions that differ significantly (Figure 5B). Gaussian Mixture Models (GMMs) were used to separate the distributions of each state into multiple Gaussians, which were visualized as violin plots using Seaborn^55^ (Figure 5C, S5B). Akaike Information Criterion and Bayesian Information Criterion scores were compared to find the optimum number of Gaussians in each distribution.

### Quantification and Statistical Analysis

All experiments were performed with at least two independent protein preparations for each DCX construct, MAP, or motor protein used. All data, unless otherwise stated, was analyzed manually using ImageJ (FIJI), with each straight and uninterrupted stretch of MT counting as a single data point. Background-subtracted mean intensities were obtained for a line scan along the MT. For kinesin-1 experiments, max projection images from 3-minute movies were generated and fluorescence intensity was quantified along microtubules. For envelope enrichment assays, mean intensities were measured inside and outside envelope boundaries as determined by WT DCX-sfGFP fluorescence. Statistical tests were performed with an unpaired, two-tailed Student’s t-test. Graphs were created using GraphPad Prism and statistical tests were performed using this program. All variances given represent standard deviation. The statistical details of each experiment can be found in the figure legends, including statistical tests used, exact value of “n”, what “n” represents and the definition of how data is plotted in each graph. Significance is defined in the figure legends: **** indicates p < 0.0001, *** indicates p < 0.001, ** indicates p < 0.01, * indicates p < 0.05, and “ns” indicates p > 0.05 where there is no statistically significant difference. When the p-value was above 0.0001, the exact p-values is reported.

