## Supplemental Figures for "Doublecortin compacts the microtubule lattice through a DC1-dependent mechanism"

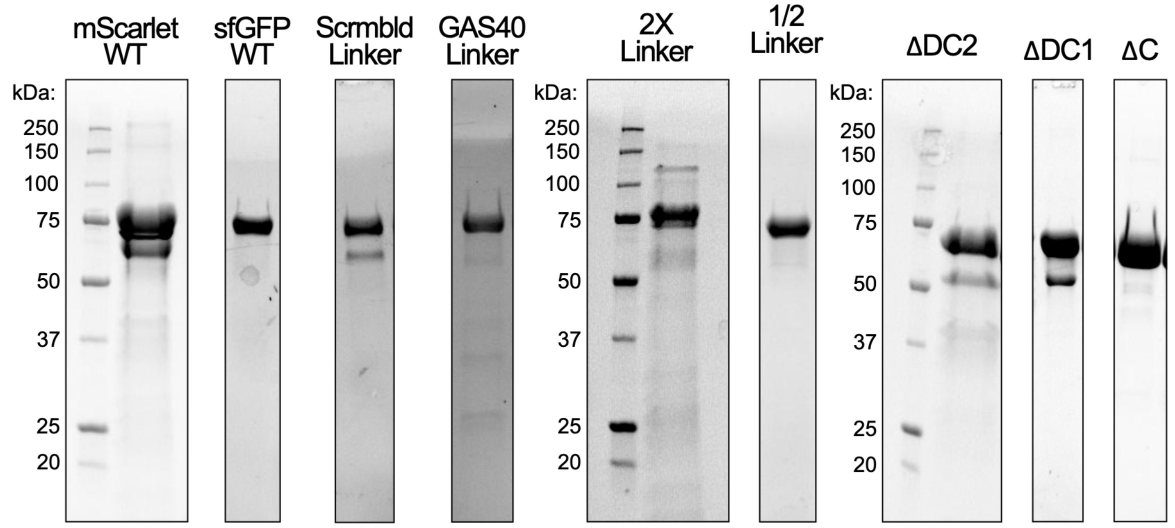


**Figure S1. Coomassie Blue-stained SDS-PAGE gels for purified recombinant proteins used in this study. Related to Figures 1-4.** Molecular weight markers (kDa) are indicated on the left.

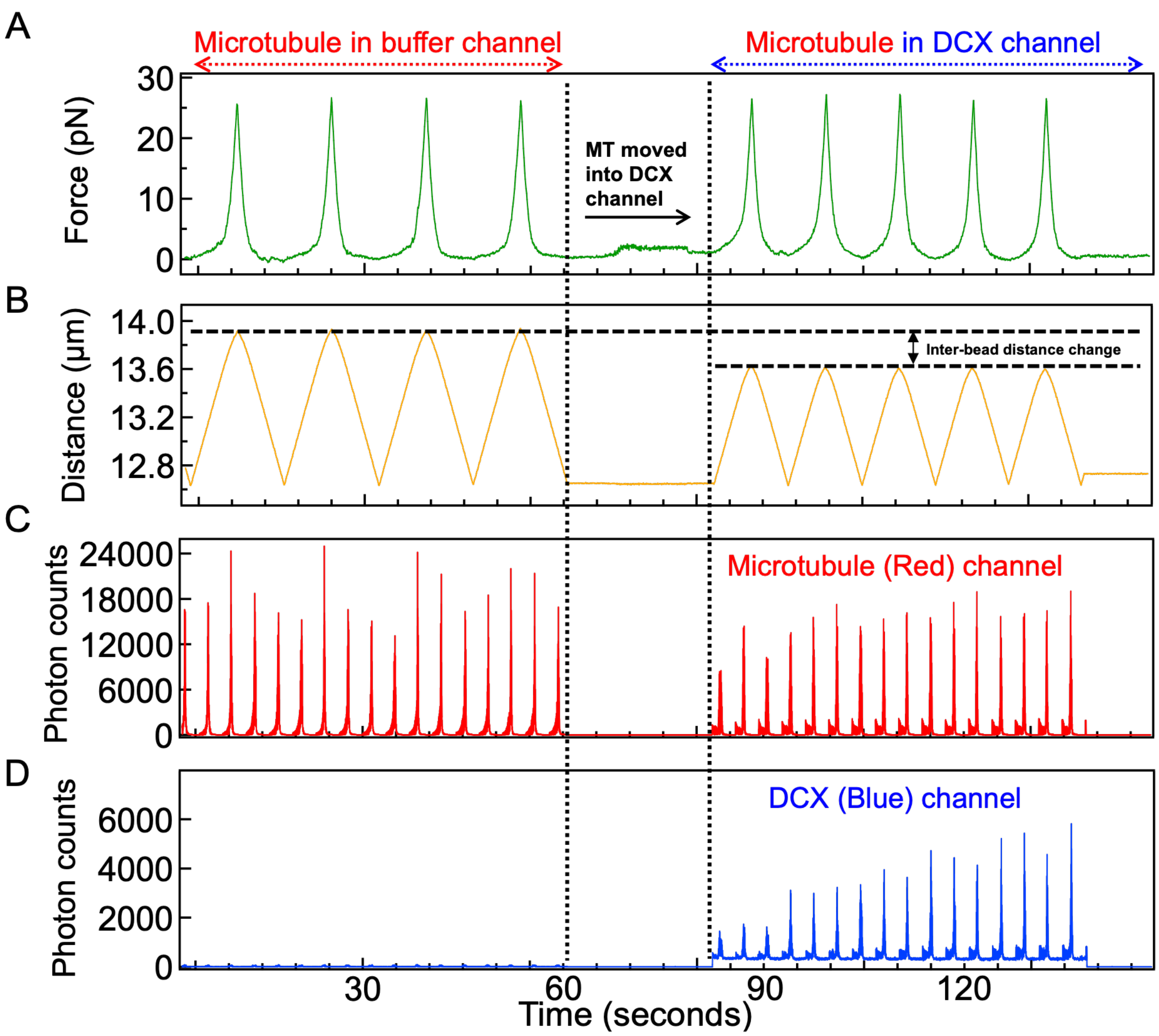


**Figure S2. DCX binding decreases microtubule inter-bead distance. Related to Figure 2.** Representative time traces show (A) force (pN), (B) inter-bead distance (µm), (C) photon counts for microtubule in the red channel under 638 nm excitation, and (D) DCX in the blue channel under 488 nm excitation. Top arrows on the time trace mark measurements done in buffer, and following the transitioning into the DCX channels across all signal windows. In panels A and B, the initial four force–extension cycles correspond to the microtubule suspended between two beads in the buffer channel. The arrow (between the vertical dashed lines) indicates the transition of the bead-microtubule tether into the DCX channel, where the five stretch-relax cycles correspond to DCX-bound microtubules. Binding of DCX to the microtubule causes a distinct change in the inter-bead distance (panel B). Consistent photon counts in the red channel confirms the presence of Alexa647-labelled microtubule throughout the experiment, while the increase in blue-channel photon counts indicates binding of sfGFP tagged DCX to the microtubules as soon as the microtubule was moved into the DCX channel.

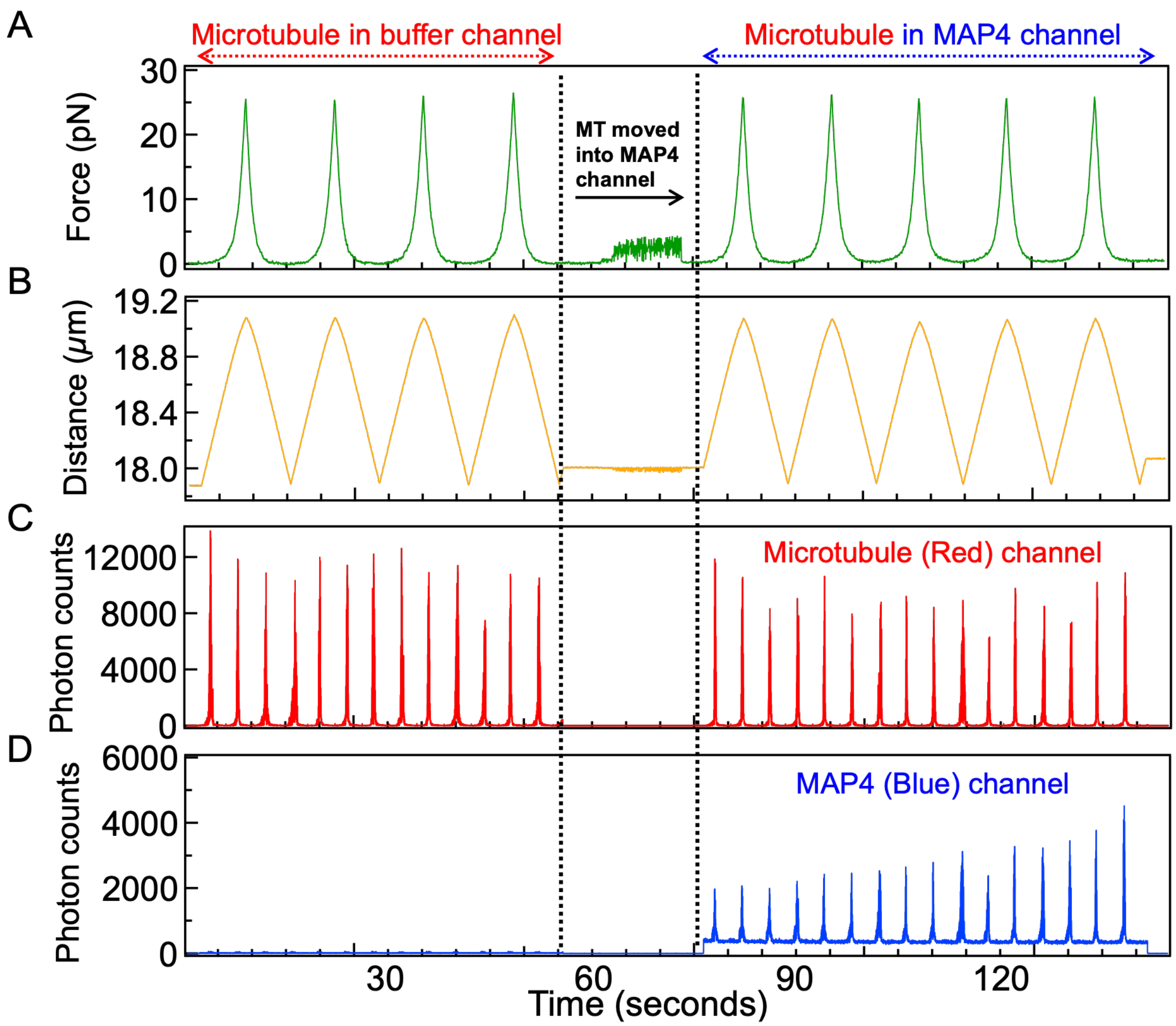


**Figure S3. MAP4 binding does not alter microtubule inter-bead distance. Related to Figure 2.** Representative time traces show (A) force (pN), (B) inter-bead distance (μm), (C) photon counts for microtubule in the red channel under 638 nm excitation, and (D) MAP4 in the blue channel under 488 nm excitation. Top arrows on the time trace mark measurements done in buffer, and following the transitioning into the DCX channels across all signal windows. In panels A and B, the initial four force–extension cycles correspond to the microtubule suspended between two beads in the buffer channel. The arrow (between the vertical dashed lines) indicates the transition of the bead-microtubule tether into the MAP4 channel, where the five stretch-relax cycles correspond to MAP4-bound microtubules. Binding of MAP4 to the microtubule did not alter the inter-bead distance (panel B). Consistent photon counts in the red channel confirms the presence of Alexa647-labelled microtubule throughout the experiment, while the increase in blue-channel photon counts indicates binding of sfGFP tagged MAP4 to the microtubules as soon as the microtubule was moved into the MAP4 channel.

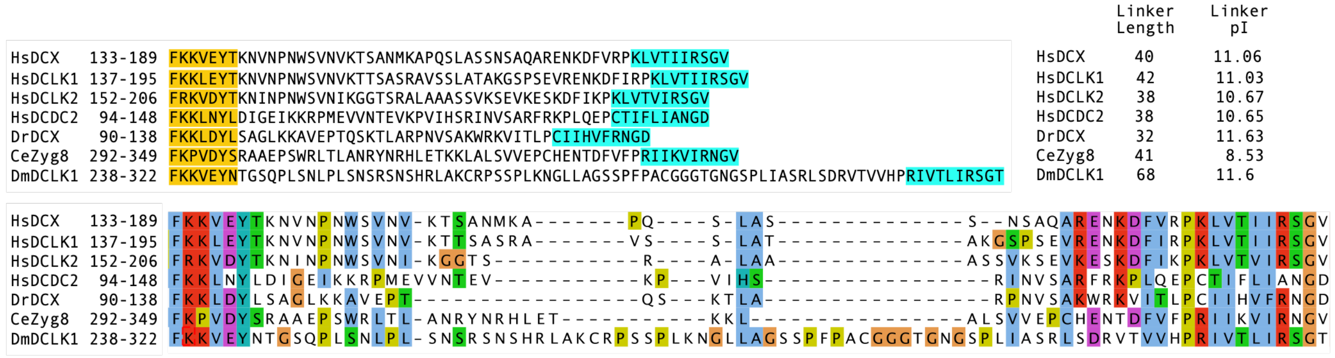


**Figure S4. Conservation of the DCX linker region. Related to Figure 3.** Top: Amino acid sequences comprising the last seven amino acids of the DC1 domain, the intervening linker region, and the first 10 amino acids of the DC2 domain are shown for each protein. The length and predicted isoelectric point (pI) of each linker region are indicated. UniProt accession number: *Homo sapiens* DCX (O43602), *Homo sapiens* DCLK1 (O15075), *Homo sapiens* DCLK2 (Q8N568), *Homo sapiens* DCDC2 (Q9UHG0), *Danio rerio* DCX (A0A8M6Z9L0), *Caenorhabditis elegans* ZYG-8 (Q95QC4), and *Drosophila melanogaster* DCLK (Q7PLI7). Bottom: The same sequences were aligned using Clustal Omega (ClustalO) and color-coded according to amino acid conservation to highlight conserved residues within the DCX linker region and flanking DCX domains.

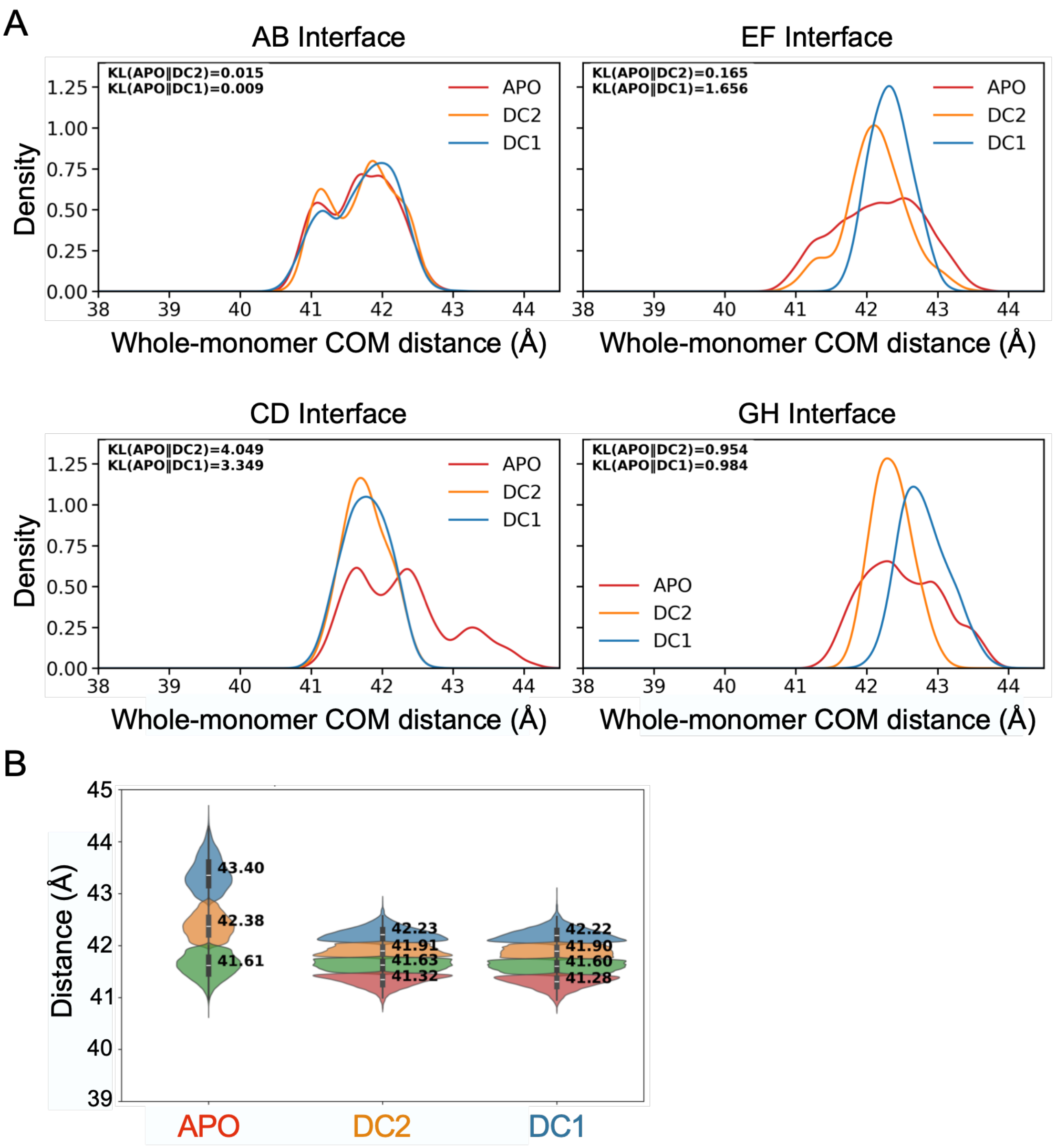


**Figure S5. Structural analyses and COM distances between monomers for APO, DC1, and DC2 systems. Related to Figure 5.** (A) Probability distributions of COM distances between interface monomer pairs for APO, DC1, and DC2. The distributions are the plus-end intradimer pairs (AB interface and EF interface) and the minus-end intradimer pairs (CD interface and GH interface). (B) Violin plots of the COM CD intradimer distances for metastable states predicted by the Gaussian Mixture Model of each system's distribution.
